# Alpha-synuclein overexpression triggers divergent cellular responses and post-translational modifications in SH-SY5Y and ReNcell VM models

**DOI:** 10.1101/2025.08.23.671356

**Authors:** Miraj Ud Din Momand, Petra Majerova, Diana Mjartinova, Natalia Maruskinova, Karolina Albertusova, Michael Dobrota, Lubica Fialova, Sara Stefankova, Petar Podlesniy, Muhammad Khalid Muhammadi, Miroslav Balaz, Dominika Fricova

## Abstract

Alpha-synuclein (α-syn) overexpression models are widely used to unravel the molecular mechanisms of Parkinson’s disease (PD), particularly in light of the dose-dependent transition between its physiological and toxic roles. However, existing systems rely on inducible expression, lack robust dose stratification and comparative cellular contexts. Here, we developed and characterized a panel of stable neuronal cell lines in two human cellular models (SH-SY5Y neuroblastoma cells and ReNcell VM neural progenitors) overexpressing GFP-tagged wild-type (WT) or A53T mutant α-syn at low and high overexpression levels. Utilizing this framework, we demonstrated that A53T consistently induces cytotoxicity, oxidative stress and mitochondrial dysfunction in both cell types. In contrast, WT α-syn had divergent effects depending on the cellular context. In SH-SY5Y cells, it enhanced mitochondrial function and viability, whereas in ReNcell VM cells, the same protein triggered mitochondrial impairment and elevated oxidative stress. This opposing metabolic response was reflected in increased respiratory activity in SH-SY5Y cells and a marked decline across WT α-syn overexpressing ReNcell VM. Importantly, post-translational modification (PTM) landscape of overexpressed WT α-syn varied dramatically by cell type. ReNcell VM cells exhibited more robust modifications signatures, even in the absence of overt aggregation, which highlights a cell-type-specific PTM landscape that may contribute to differential vulnerability. Our findings underscore a complex interplay between α-syn dosage, mutational status, cellular environment, and PTM profiles highlighting that neuronal vulnerability in PD is context-dependent. This work establishes a modular in vitro platform for dissecting α-syn pathology and testing targeted therapeutic strategies grounded in cell-type specificity.

## Introduction

Cell-based models remain essential for investigating alpha-synuclein (α-syn)-related pathology in Parkinson’s disease (PD), as they allow controlled manipulation of expression levels and capture the dose-dependent shift in α-syn function from physiological to pathological states. While physiological levels support normal neuronal activity, elevated expression as observed in gene multiplications linked to familial PD [1–3], drives toxicity through mechanisms such as mitochondrial dysfunction, impaired vesicle trafficking, and altered protein homeostasis [4, 5]. These models have employed diverse cellular backgrounds, ranging from non-neuronal HEK293T and H4 neuroglioma cells [6] to human-derived neuronal models such as SH-SY5Y neuroblastoma cells [7, 8], primary neurons [8], patient-derived fibroblasts, and induced pluripotent stem cells [9]. Among these models, SH-SY5Y neuroblastoma cells stably overexpressing wild-type (WT) or PD-related A53T mutant form of α-syn have emerged as a cornerstone for studying α-syn-mediated toxicity. Their widespread use has provided mechanistic insights into several key aspects of PD pathology. Notably, α-syn overexpression in SH-SY5Y cells has been shown to induce marked mitochondrial fragmentation, primarily through inhibition of mitochondrial fusion and promotion of fission processes, with A53T-expressing cells exhibiting more pronounced phenotypes [10]. α-syn overexpression in SH-SY5Y revealed impaired calcium homeostasis, particularly at mitochondria-associated endoplasmic reticulum membranes, where α-syn disrupts calcium transfer [11]. Further studies using SH-SY5Y cells have demonstrated that elevated α-syn impairs autophagic flux and lysosomal function, contributing to defective clearance of protein aggregates [12]. Additionally, α-syn-overexpressing SH-SY5Y cells display dysregulated iron metabolism, including altered iron uptake and homeostasis, which may contribute to oxidative stress [13]. Exosomes derived from α-syn overexpressing SH-SY5Y cells have been shown to cause impairment of autophagy in recipient cells [14]. Beyond functional impairments, these models have been instrumental in the characterization of α-syn post-translational modifications (PTMs) related to PD, especially phosphorylation, which is commonly observed in Lewy body inclusions in PD brains [15–17]. Despite their utility, SH-SY5Y cells carry limitations that constrain their physiological relevance. They represent a single cancer-derived lineage, often lack stratified expression models to test dosage-dependent effects, and have not been extensively compared to other human neuronal models.

In this study, we established a robust dual-cell model to address these limitations by using two human-derived neuronal lines; SH-SY5Y and ReNcell VM. ReNcell VM, originating from human fetal ventral mesencephalon and immortalized via v-myc, more closely resembles neural progenitors of the substantia nigra, a region critically affected in PD, making it a highly relevant model for comparison. We introduced stable overexpression of GFP-tagged WT or A53T α-syn via lentiviral vectors in both cell types and stratified the populations into low and high α-syn expressers, enabling dose-dependent comparisons. This multifaceted approach allowed us to directly compare α-syn-induced effects in two distinct neuronal cell types, investigate how WT and mutant A53T α-syn differentially impact cellular homeostasis, systematically assess dose-dependent responses to α-syn overload, a rarely addressed dimension in previous models. Our findings reveal that ReNcell VM cells exhibit pronounced sensitivity to α-syn-induced mitochondrial dysfunction and display a distinct PTM profile compared to SH-SY5Y cells. While previous models using SH-SY5Y cells have highlighted phosphorylation at serine-129 (pS129) as a hallmark of α-syn pathology, our study demonstrates that this modification occurs more robustly and in a broader PTM landscape in ReNcell VM cells, even in the absence of overt aggregation. This suggests that specific cellular environments can influence conformational state and pathogenic transition of α-syn. Collectively, these findings underscore the critical importance of cellular context, α-syn variant, and overexpression dosage in shaping pathological outcomes, and they highlight the need for diversified models to more accurately capture early disease mechanisms in PD.

## Materials and Methods

### Generation and maintenance of stable cell lines

Two cell lines used in this study were human neuroblastoma SH-SY5Y cells (Sigma-Aldrich, USA, 94030304) and human neural progenitor cell line ReNcell VM (Millipore, Germany, SCC008), generated by insertion of a v-myc oncogene into human fetal ventral mesencephalic cells. SH-SY5Y cells were maintained in DMEM medium (Gibco, USA, 11960044) supplemented with 10% FBS (Biosera, France, FB-1090/500), 2 mM L-glutamine (Biosera, France, XC-T1715/100), 1% penicillin/streptomycin (100 U/ml penicillin, 100 µg/ml streptomycin) (Biosera, France, LM-A4118). ReNcell VM cells were maintained in Matrigel (Corning, USA, 356234) coated plates with ReNcell NSC Maintenance Medium (Sigma-Aldrich, USA, SCM005) supplemented with 20 ng/ml fibroblast growth factor-2, 20 ng/ml epidermal growth factor, 1% penicillin/streptomycin. Cell lines stably overexpressing WT or A53T mutant α-syn were generated using eGFP-α-syn (referred to in text as GFP-α-syn) WT or A53T plasmids, a gift from David Rubinsztein (via Addgene, USA, plasmids 40822 or 40823) through lentivirus transduction.

### Cell sorting and flow cytometry analysis

Stably transduced cells were sorted based on GFP expression using CytoFLEX SRT (Beckman, USA). Cells were resuspended in phosphate-buffered saline (PBS) for sorting, GFP was excited at 488 nm, and emission was collected using a 525/40 nm filter. Cell lines were sorted into low and high overexpression populations. Sorted cells were collected in a tube containing growth medium and transferred to cell culture dishes. Successful sorting of GFP-positive cells was confirmed through microscopy. These cells were then expanded and used for subsequent experiments. To minimize variability and potential adaptation to α-syn-induced stress, all cell lines were either freshly thawed or maintained in culture for no more than five passages. For CellROX assay, cells were seeded and stained upon reaching 70% confluence with CellROX Orange reagent (Thermo Scientific, USA, C10443) at a final concentration of 2.5 µM for 30 min in growth conditions (humidified chamber with 37°C, 5% CO₂). After 30 min, cells were detached, resuspended in PBS and analyzed via flow cytometry using LSRFortessa (BD Biosciences, USA) with 488 nm laser and 540/40 nm filter for GFP and 561 nm laser and 586/15 nm filter for CellROX Orange. For mitochondrial staining cells were stained with 1 µM MitoTracker Red (Thermo Scientific, USA, M22426) solution for 30 min in growth conditions. After incubation and detachment, cells were resuspended in 500 µl PBS supplemented with 5% FBS. Cell suspensions were analyzed on a MACSQuant (Miltenyi Biotec, Germany) flow cytometry analyzer using a 488 nm laser and a 585/40 nm filter. 10,000 cell events were probed and compared to unstained control cells. Data acquisition and analysis were performed using FCS Express 7 software (De Novo software, USA).

### Quantitative Real-Time Polymerase Chain Reaction (qPCR)

Total RNA was extracted from cells using High Pure RNA Isolation Kit (Roche, Switzerland, 11828665001). cDNA was synthesized with 100 ng of total RNA using High-Capacity cDNA Reverse Transcription Kit (Applied Biosystems, USA, 4368814). Diluted cDNA (1:10) was used for gene expression analysis, each sample in triplicates mixed with 1x TaqMan Gene Expression Master Mix and relevant 1x TaqMan probes (Applied Biosystems, USA) Hs00240906_m1 or Hs04194366_g1 targeting SNCA or housekeeping gene (RPL13A) respectively. qPCR was performed on a QuantStudio 5 RT-PCR System (Applied Biosystems, USA). Reactions (10 µl) were performed using the following cycling program: hold at 50°C for 2 min, initial denaturation at 95°C for 10 min, followed by 40 cycles of denaturation for 15 seconds and annealing/extension at 60°C for 1 minute. Relative mRNA levels were calculated according to the 2−ΔΔ*C*_t_ method [18].

### Immunoblotting

Cells were lysed in RIPA or M-PER Mammalian Protein Extraction Reagent (Thermo Scientific, USA, 78503) supplemented with cOmplete protease inhibitor cocktail (Roche, Switzerland, 05056489001) on ice for 30 min, centrifuged at 13,000xg for 15 min and quantified with Pierce BCA Protein Assay Kit (Thermo Scientific, USA, 23225). Equal amounts of protein were separated on SDS-PAGE and transferred to a nitrocellulose (Cytiva, USA, 45004001) or polyvinylidene fluoride (PVDF; Thermo Scientific, USA, 88518) membrane. For α-syn detection, the membranes were fixed with 0.4% paraformaldehyde (PFA) and then blocked with 5% milk. The membranes were probed overnight with the antibodies Syn211 (Invitrogen, USA, AHB0261), HSP90 (Cell Signaling Technology, USA, 4877), anti phospho-α-syn pS129 (Cell Signaling Technology, USA, 23706), GAPDH (Cell Signaling Technology, 5174) diluted 1:1000, and anti-GFP tag antibody (Proteintech, USA, 66002-1-Ig) diluted 1:5000. Membranes were then incubated with suitable secondary antibodies conjugated to horseradish peroxidase (Agilent, USA, P0447, P0448). The signal was detected using SuperSignal West Pico PLUS Chemiluminescent Substrate (Thermo Scientific, USA, 34580) on iBright 1500 Imaging System (Invitrogen, USA).

### Immunocytochemical and mitochondrial staining

For immunocytochemistry staining cells were washed with PBS, fixed in 4% PFA for 15 min, and permeabilized with 0.1% Triton X-100 for 30 min at room temperature. Non-specific binding was blocked with 2% BSA (Sigma-Aldrich, USA, 9048-46-8) for 1 h at room temperature and stained with primary antibody (Syn211, Invitrogene, 32-8100; pS129, Cell Signalling Technology, 23706) diluted 1:500. All primary antibodies were diluted in 0.1% BSA and incubated overnight at 4°C. Cells were subsequently incubated with compatible secondary antibodies conjugated to Alexa-546 or Alexa-647 fluorophores (Invitrogen, USA, A-11030, A-21245) diluted 1:1000 for 1 h at room temperature. For mitochondrial staining cells were stained with 1 µM MitoTracker Red (Thermo Scientific, USA, M22426) dye solution for 30 min in growth conditions. Subsequently, cells were washed with 1x PBS and fixed with 4% PFA at room temperature for 10 min. The nuclei were stained with Hoechst (Invitrogen, USA, 62249) for 10 min, and cells were mounted in Fluoromount-G mounting medium (Merck, USA, F4680). Stained samples were imaged with Zeiss LSM 710 confocal microscope (Zeiss, Germany).

### Cell viability analysis

Both SH-SY5Y and ReNcell VM cell lines were seeded in respective growth conditions in triplicates at a starting density of 10000 cells/well in 96 well plates. Cell viability was assessed after cells reached around 70% confluence by adding MTT labeling reagent (Roche, Switzerland, 11465007001) with final concentration 0.5 mg/ml to each well and cells were incubated for 4 h in growth conditions. After 4 h, 100 µl of the solubilization buffer was added into each well and the plate was incubated overnight in growth conditions. The following day absorbance was measured with Varioskan plate reader (Thermo Scientific, USA) at wavelengths of 570 nm and 690 nm. For In Vitro Toxicology Resazurin-based assay (Merck, USA, TOX8) cells were seeded in a similar way. After reaching 70% confluence, resazurin dye solution was added to each well at an amount corresponding to 10% of the total well volume. Cells were incubated for 3 h in growth conditions and fluorescence was measured using a Varioskan multiplate reader with excitation at 560 nm and emission at 590 nm.

### Immunoprecipitation and liquid chromatography with mass spectrometry (LC/MS) analysis

GFP-α-syn was immunoprecipitated using ChromoTek GFP-Trap Magnetic Agarose beads (Proteintech, USA). Briefly, 300 µg of total protein lysates were incubated overnight with 25 µl of beads on DYNAL Sample Mixer (Dynal Biotech, Norway) at 4°C. The following day, beads were spun at 800 rpm for 30 seconds, supernatant was collected, beads were washed once, and the proteins were eluted in 100 µl of 0.2 M glycine, pH 2.5. pH was neutralized with Tris, pH 9.0 and successful pull-down was validated through immunoblotting with anti-GFP tag antibody. For LC/MS analysis, samples were dried using a SpeedVac and diluted in 8 M urea. Reduction was performed with 10 mM dithiothreitol (Sigma-Aldrich, USA, D9779) at 37°C for 1 h. Alkylation was carried out with 15 mM iodoacetamide (Sigma-Aldrich, USA, I1149), protected from light for 30 min. Trypsin digestion (Promega, USA, V5280) was performed overnight at 37°C. The resulting peptide mixtures were separated using an Acquity M-Class Ultra-High-Performance Liquid Chromatography system (Waters, USA), employing a nanoEase Symmetry C18 trap column (20 mm length, 180 µm diameter, 5 µm particle size; Waters, USA) for desalting, followed by separation on a nanoEase High Strength Silica T3 C18 analytical column (100 mm length, 75 µm diameter, 1.8 µm particle size). A 75-minute gradient of acetonitrile (5% to 40%) (Sigma-Aldrich, USA, 45983) containing 0.1% formic acid (Sigma-Aldrich, USA, 5.33002) was applied at a flow rate of 300 nL/min. Analysis was conducted using a Synapt G2-Si quadrupole time-of-flight mass spectrometer with ion mobility (Waters, USA). Data was acquired in a data-independent acquisition (DIA) mode at a spectral acquisition rate of one second. Peak detection and data processing were performed using PEAKS Studio 12.5 (Bioinformatics Solutions, Canada).

### Mitochondrial bioenergetics profiling using Seahorse XF96

Mitochondrial respiration was assessed using the Seahorse XF96 Extracellular Flux Analyzer (Agilent, USA). SH-SY5Y and ReNcell VM cell lines were seeded at a density of 30,000 cells per well in Seahorse XF96 cell culture microplates (Agilent, USA, 103794-100) coated with Matrigel and incubated overnight in standard growth conditions. The sensor cartridge was hydrated in Seahorse XF Calibrant (Agilent, USA, 100840-000) and placed in a 37°C non-CO_2_ incubator overnight. On the day of the assay, the culture medium was replaced with Seahorse XF DMEM assay medium (Agilent, USA, 103680-100) supplemented with 10 mM glucose (Agilent, USA, 103577-100), 1 mM pyruvate (Agilent, USA, 103578-100), and 2 mM glutamine (Agilent, USA, 103579-100), adjusted to pH 7.4. Cells were incubated at 37°C in a non-CO_2_ incubator for 60 min. The compounds were diluted in complete Seahorse XF DMEM medium and loaded into the appropriate ports of the sensor cartridge to reach final concentrations of 1.5 µM oligomycin, 2 µM FCCP, and 0.5 µM rotenone/antimycin A. The cartridge was then placed in the Seahorse XF96 Analyzer for calibration for 20 min. Subsequently, the sensor cartridge was replaced with the cell culture plate and measurement of oxygen consumption rate (OCR) was performed. OCR was analyzed using Wave software (Agilent Technologies, Inc, USA). Data were normalized to total protein content using DC assay.

### Statistical analysis

All quantitative data are expressed as mean ± standard error of the mean (SEM), geometric mean or median, as indicated. Each experiment was independently repeated three times. Statistical significance was determined using unpaired two-tailed t test, one-way ANOVA with Dunnett’s correction, or Kruskal–Wallis test with Dunn’s post hoc test, as appropriate. Values of p < 0.03 (*), p < 0.002 (**), and p < 0.001(***) were considered statistically significant; p > 0.12 was considered not significant (ns). All statistical analyses were performed using GraphPad Prism version 10.5.0 (774). All figures were created using GraphPad Prism and Adobe Illustrator 2023.

## Results

### Generated stable α-syn overexpression neuronal cell models in SH-SY5Y and ReNcell VM cell lines exhibit consistent α-syn (WT/A53T) overexpression

To systematically investigate α-syn variant’s and dosage-dependent effects across distinct neuronal backgrounds, we developed a comprehensive panel of neuronal cell lines overexpressing α-syn through lentiviral transduction and fluorescence-activated cell sorting (FACS). Lentiviral constructs encoding either WT or A53T mutant α-syn fused to GFP leading to N-terminal fusion were generated and used to transduce two human neuronal cell lines: SH-SY5Y neuroblastoma cells and ReNcell VM ventral midbrain neural stem cells (Fig. 1A). Following transduction, cells were sorted by flow cytometry based on GFP fluorescence intensity, enabling precise selection of populations with defined overexpression levels designated as low (L) and high (H) overexpressers. Flow cytometric analysis demonstrated successful transduction efficiency in both cell lines, with clear separation between untransduced and transduced populations based on GFP fluorescence intensity. The transduced populations displayed distributions enabling effective gating strategies for low and high overexpression level selection in both SH-SY5Y and ReNcell VM cells. Untransduced control populations exhibited minimal background fluorescence, confirming the specificity of the sorting methodology (Fig. 1B). This strategy yielded eight genetically defined α-syn overexpressing cell lines: SH-SY5Y (α-syn WT L, α-syn WT H, α-syn A53T L, α-syn A53T H) and ReNcell VM (α-syn WT L, α-syn WT H, α-syn A53T L, α-syn A53T H), complemented by untransduced control lines for each cell type. qPCR analysis confirmed robust overexpression of SNCA mRNA across all generated cell lines (Fig. 1C). Both SH-SY5Y and ReNcell VM lines overexpressing WT and A53T α-syn showed significant increases in SNCA transcript levels compared to their respective controls. Notably, high-expressing groups consistently exhibited several-fold greater mRNA abundance than their low-expressing counterparts, validating the success of the GFP-based sorting strategy. However, we detected variability in SNCA mRNA expression among biological replicates that had undergone extended passaging, likely due to passage-related changes in lentiviral integration efficiency or transcript regulation. To minimize these inconsistencies, all subsequent experiments were performed using freshly thawed cells and limited to early passage numbers. Thus, despite this transcriptional variability, total α-syn protein levels remained consistent across experimental replicates, as confirmed by immunofluorescence (Fig. 3) and immunoblotting analysis (Fig. 6).

**Figure 1.**
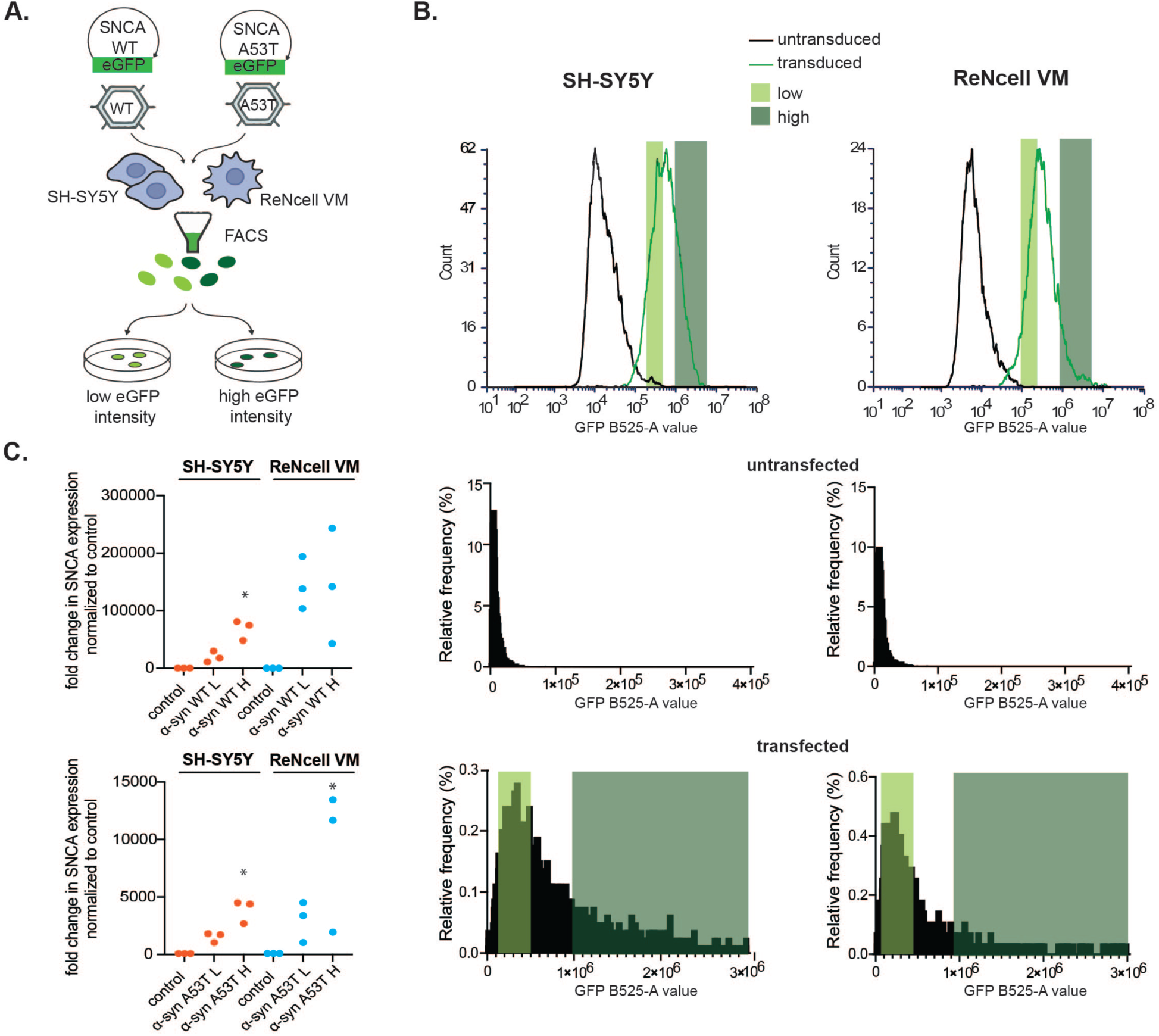
Generated stable α-syn overexpression neuronal cell models in SH-SY5Y and ReNcell VM cell lines exhibit consistent α-syn (WT/A53T) overexpression (A) Brief schematic overview of the workflow used to generate stable cell lines in SH-SY5Y/ReNcell VM cells overexpressing GFP tagged α-syn WT or A53T mutant via lentivirus transduction. Transduced cells were selected based on GFP expression with FACS. (B) Representative FACS plots illustrating the gating strategy adopted to separate GFP positive and GFP negative populations. GFP positive cells were further separated into low and high overexpression distinguished based on the fluorescent intensity of GFP. (C) qPCR analysis of SNCA mRNA expression levels in sorted cell lines in comparison with non-transduced SH-SY5Y/ReNcell VM cells, normalized to housekeeping gene RPL13A and shown here as fold change relative to the control. Dots represent three independent biological replicates, each involving separate passages, cell seeding and RNA isolations. Statistical significance was determined using an unpaired two-tailed t-test, significance indicated as p < 0.03 (*), p < 0.002 (**), p < 0.001 (***); non-significant p > 0.12).

### α-syn overexpression differentially affects cell viability and reactive oxygen species production in neuronal cell models

To assess the cellular effects of α-syn overexpression, we employed two complementary viability assays with distinct mechanistic principles: Resazurin reduction and MTT metabolism (Figure 2A, B). The resazurin assay measures overall cellular metabolic activity through the reduction of resazurin to resorufin by metabolically active cells, reflecting general cellular health and metabolic capacity. In contrast, the MTT assay specifically evaluates mitochondrial metabolic function through the reduction of tetrazolium salt by mitochondrial dehydrogenases, providing targeted assessment of mitochondrial integrity and oxidative metabolism. The resazurin assay revealed striking cell type-specific and mutation-dependent responses. SH-SY5Y cells demonstrated beneficial metabolic effects with WT α-syn overexpression, with both low and high levels showing increased metabolic activity, reaching approximately 120-150% of control values. Conversely, the A53T variants exhibited opposite effects, displaying reduced cellular viability in both the low and high overexpression groups, indicating that the pathogenic mutation transforms α-syn from metabolically beneficial to detrimental in this cellular context. ReNcell VM cells maintained viability levels comparable to controls across all α-syn overexpression conditions in the resazurin assay, indicating minimal impact on overall cellular metabolic capacity regardless of variant type or overexpression level (Figure 2A). The MTT assay provided contrasting results, highlighting the assay-specific sensitivity to α-syn overexpression and revealing differential mitochondrial vulnerability profiles. SH-SY5Y cells showed mitochondrial resilience, with no significant alterations in MTT reduction capacity across all α-syn variants and overexpression levels, maintaining mitochondrial function similar to controls. However, ReNcell VM cells demonstrated pronounced mitochondrial dysfunction, with reduced MTT metabolism observed across all α-syn overexpressing populations. Both WT and A53T variants at all overexpression levels showed decreased mitochondrial metabolic capacity compared to controls, indicating that ReNcell VM cells are inherently more susceptible to α-syn-induced mitochondrial impairment (Figure 2B).

**Figure 2.**
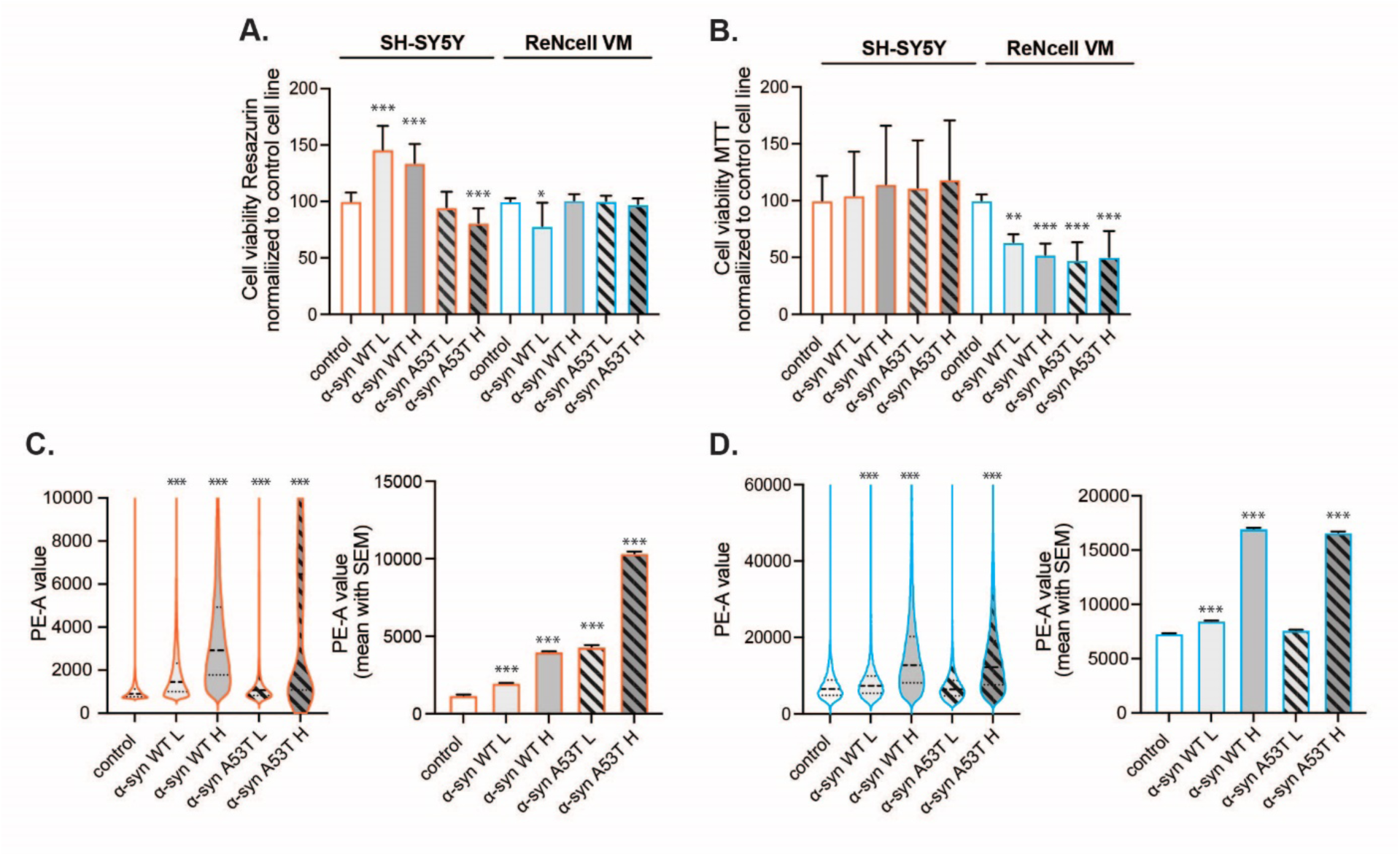
α-syn overexpression differentially affects cell viability and ROS production in neuronal cell models. The effects of α-syn (WT/A53T) overexpression on cell viability were assessed using (A) resazurin and (B) MTT assays in SH-SY5Y and ReNcell VM cell lines. Viability is presented here as a percentage relative to untransduced control cells. Group comparisons were analyzed by one-way ANOVA followed by Dunnett’s correction for multiple comparisons. (C–D) Intracellular reactive oxygen species (ROS) production was measured in α-syn (WT/A53T) overexpressing SH-SY5Y (C) and ReNcell VM (D) cells using CellROX Orange staining followed by flow cytometry revealing elevated ROS levels compared to untransduced controls. Violin plots and corresponding bar graphs summarizing mean fluorescence intensity are shown displaying the distribution/shift and mean ± SEM of CellROX fluorescence intensity. Group comparisons were performed using the Kruskal–Wallis test with Dunn’s post hoc correction for multiple comparisons. Statistical significance is indicated as p < 0.03 (*), p < 0.002 (**), p < 0.001 (***); non-significant p > 0.12).

Reactive oxygen species analysis using CellROX staining and flow cytometric quantification revealed distinct cellular oxidative stress responses to α-syn overexpression (Figure 2C, D). In SH-SY5Y cells, reactive oxygen species (ROS) production demonstrated a graduated mutation-dependent pattern, with substantial progressive increases from WT low overexpression through A53T high overexpression conditions. The most pronounced ROS elevation occurred in A53T high overexpressers, indicating that both mutation status and overexpression level contribute to oxidative burden in these cells (Figure 2C). ReNcell VM cells exhibited broader sensitivity to α-syn-induced oxidative stress, with both WT and A53T variants inducing elevated ROS production compared to controls. The oxidative stress response was most prominent in high overexpression conditions regardless of variant type. Notably, ReNcell VM cells demonstrated substantially higher ROS production responses compared to SH-SY5Y cells across comparable overexpression conditions, indicating enhanced susceptibility to α-syn-induced oxidative stress (Figure 2D).

### α**-**syn localization and phosphorylation are variant and dose-dependent but do not differ between the neuronal cell models

To assess the subcellular distribution and pathological modification of α-syn in our cellular models, we performed immunocytochemistry using conformation-specific Syn211 antibody and phospho-specific anti-pS129 antibody across SH-SY5Y and ReNcell VM backgrounds, with variable overexpression levels of WT or A53T α-syn (Fig. 3). Immunofluorescence microscopy utilizing the Syn211 antibody (Fig. 3A, B), which specifically recognizes amino acid residues 121-125 of α-syn, validated successful overexpression of GFP-α-syn fusion constructs across all generated cell lines. The analysis demonstrated robust colocalization between GFP fluorescence and Syn211 immunoreactivity in both SH-SY5Y and ReNcell VM cells, confirming fusion protein integrity and detectability. Control cells exhibited minimal staining with the Syn211 antibody, establishing the specificity of α-syn detection. Quantification of Syn211 signal intensity relative to control cells confirmed a marked increase in α-syn protein levels across all overexpressing lines (Fig. 3E). Both SH-SY5Y and ReNcell VM cell lines overexpressing WT α-syn demonstrated a predominantly diffuse, homogeneous cytoplasmic distribution pattern independent of overexpression levels (Fig. 3A, B). Low expression WT cells displayed evenly distributed fluorescence throughout the cytoplasm. High overexpression WT variants maintained this diffuse localization pattern, exhibiting increased fluorescence intensity without formation of discrete intracellular aggregates or punctate structures. Cells overexpressing the A53T pathogenic mutation exhibited markedly distinct subcellular localization characterized by formation of prominent punctate structures. Both low and high overexpression A53T variants demonstrated cytoplasmic inclusions with aggregate-like morphology across both cell lines (Fig. 3A, B). These punctate structures appeared as discrete, intensely fluorescent foci distributed throughout the cytoplasm, readily distinguishable from the diffuse pattern observed in WT overexpressing cells. Immunofluorescence analysis utilizing pS129-specific antibodies revealed modification patterns that closely mirrored the subcellular distribution observed with Syn211 staining (Fig. 3C, D). Phosphorylated α-syn at serine 129 represents a well-characterized pathological marker in PD and synucleinopathies. WT α-syn overexpressing cells in both SH-SY5Y and ReNcell VM backgrounds demonstrated diffuse cytoplasmic pS129 immunoreactivity corresponding to the homogeneous distribution pattern observed with total α-syn detection. Low overexpression WT groups exhibited modest pS129 signal intensity, while high overexpression WT variants showed proportionally increased pS129 staining intensity throughout the cytoplasm, maintaining the diffuse localization pattern without formation of discrete phosphorylated aggregates (Fig 3A-D). Cells overexpressing A53T mutant α-syn exhibited markedly enhanced pS129 immunoreactivity with a distinct punctate distribution pattern (Fig. 3C, D). Both low and high overexpression A53T variants demonstrated intense pS129 staining that strongly co-localized with the aggregate-like punctate structures observed in total α-syn detection. The pS129 modification patterns remained consistent across both SH-SY5Y and ReNcell VM cell types, with no discernible differences in staining intensity or distribution between the neuronal backgrounds

**Figure 3.**
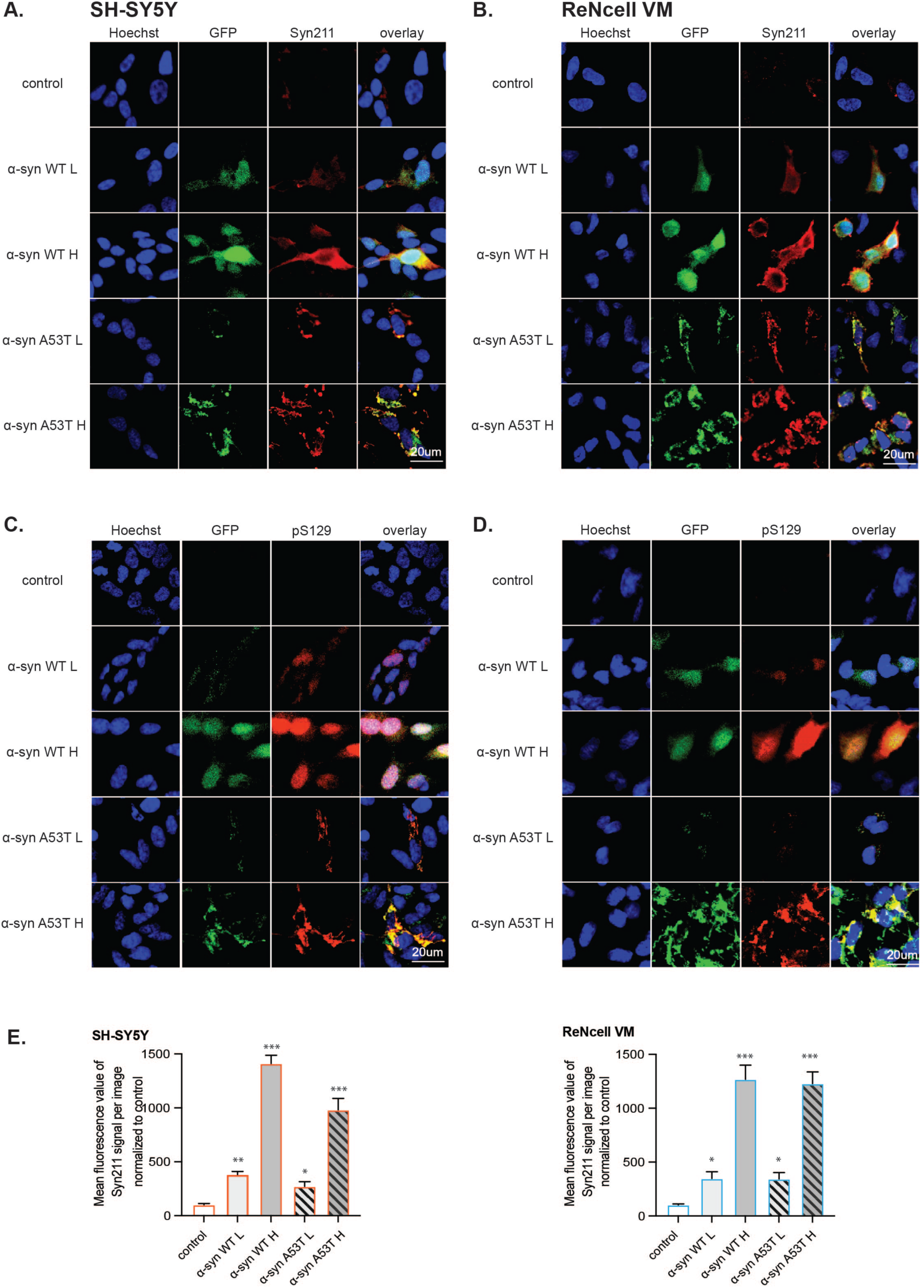
α-syn localization and phosphorylation are variant- and dose-dependent but do not differ between the neuronal cell models. Immunocytochemistry staining was performed to assess the subcellular distribution and pathological modification status of WT/A53T α-syn in SH-SY5Y and ReNcell VM cell lines. Total α-syn was detected using the Syn211 antibody in (A) SH-SY5Y and (B) ReNcell VM cells that robustly colocalized with GFP, while phosphorylated α-syn at serine 129 (pS129) was visualized using phospho-specific antibody pS129 in SH-SY5Y (C) and ReNcell VM (D) cells. Overexpression-level and variant-specific differences in subcellular distribution were observed, with A53T higher overexpression associated with more intense and distinct punctate signals. Nuclei were counterstained with Hoechst. (E) Microscopic signal intensity was quantified using ImageJ software by measuring the mean fluorescence intensity within individual cells, with background subtraction applied uniformly across images. At least 10 cells per condition were analyzed, and values were normalized to the mean intensity of non-overexpressing control cells to assess relative α-syn protein overexpression levels. Group comparisons were analyzed using one-way ANOVA with Dunnett’s correction for multiple comparisons with significance indicated as p < 0.03 (*), p < 0.002 (**), p < 0.001 (***); non-significant p > 0.12).

### α-syn overexpression alters mitochondrial integrity in a variant-, dose- and cell-type dependent manner in neuronal cell models

Following our observation of cell type-specific mitochondrial dysfunction in MTT assays, we investigated the potential of the underlying mitochondrial alterations induced by α-syn overexpression using MitoTracker Red staining and quantitative flow cytometric analysis across generated SH-SY5Y and ReNcell VM cell line panel (Fig. 4). MitoTracker Red dye accumulates in metabolically active mitochondria with intact membrane potentials, providing quantitative assessment of mitochondrial mass and functional status. This analysis revealed profound cell type-specific and variant-dependent differences in mitochondrial properties between the neuronal models. In SH-SY5Y cells we observed in both low and high WT α-syn overexpression groups typical elongated and tubular mitochondrial morphology and significantly increased MitoTracker Red fluorescence intensity compared to control (Fig. 4A). These cell models exhibited quantitatively enhanced mitochondrial staining (Fig. 4A), suggesting either increased mitochondrial mass or enhanced mitochondrial membrane potential. SH-SY5Y cells overexpressing A53T mutant α-syn displayed distinct mitochondrial morphology changes. Mitochondria appeared more punctate and condensed suggesting increased mitochondrial fragmentation (Fig. 4A). This was further supported by quantification, where the increased MitoTracker Red signal reflected an increase in mitochondrial density (Fig. 4C). MitoTracker Red staining was weaker in the ReNcells VM compared to SH-SY5Y (Fig. 4B). In ReNcell VM cells we did not observe morphological changes between WT α-syn overexpressing cells and control, but quantification of MitoTracker Red signal showed decreased fluorescence intensity compared to controls across both low and high overexpression groups (Fig. 4D). This reduction in mitochondrial staining suggests either decreased mitochondrial mass or reduced mitochondrial membrane potential, indicating mitochondrial dysfunction in this cellular context. ReNcell VM cells highly overexpressing A53T mutant α-syn demonstrated analogous alterations in mitochondrial architecture as observed in SH-SY5Y cell lines and a shift toward fragmented and granular pattern (Fig. 4B). High-overexpression conditions of A53T resulted in extensive mitochondrial clustering, and loss of filamentous structure. These morphological disruptions were quantitatively confirmed by significant increases in mitochondrial signal density, especially in the A53T H group (Fig. 4D).

**Figure 4.**
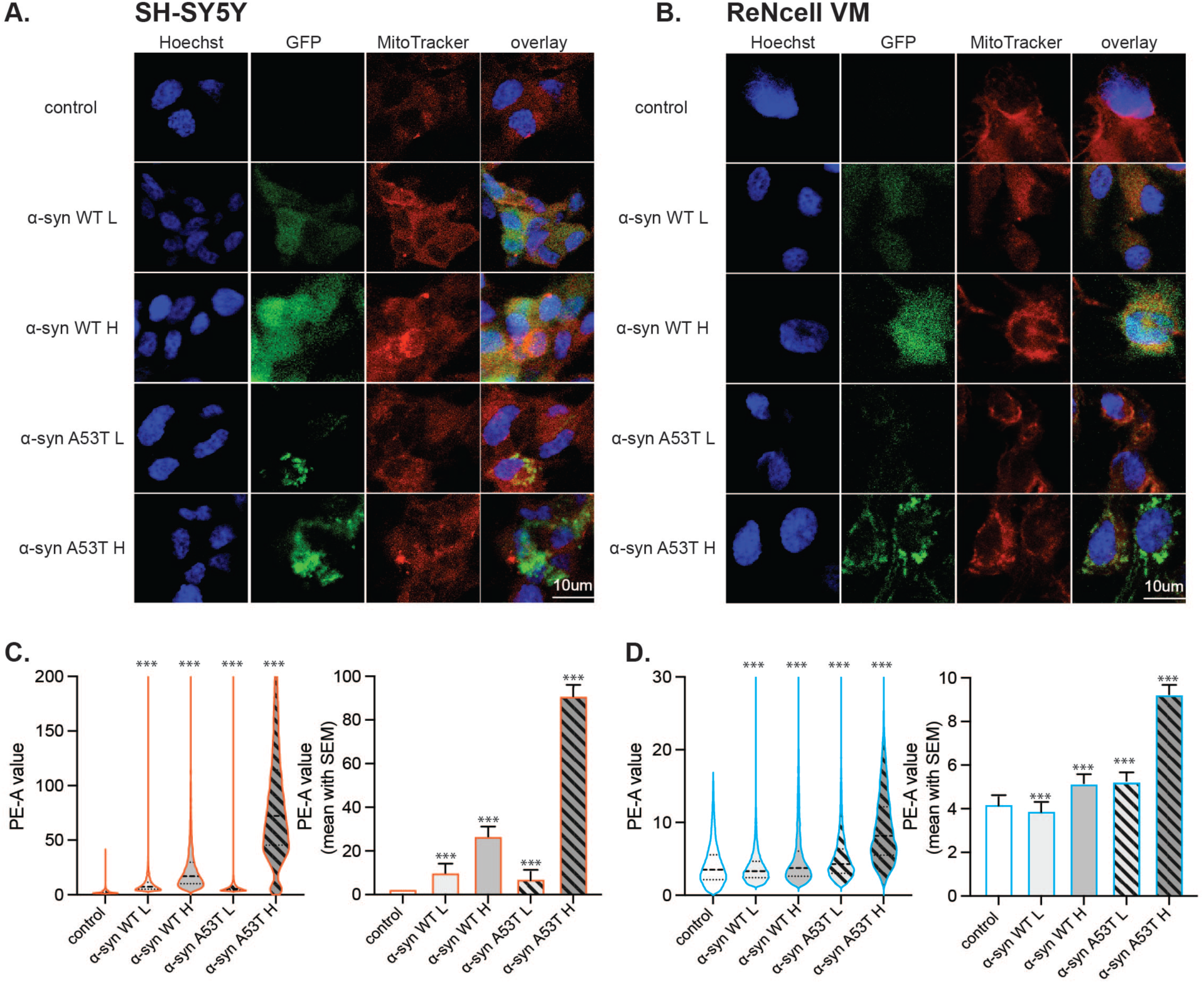
α-syn overexpression alters mitochondrial integrity in a variant-, dose- and cell-type dependent manner in neuronal cell models. Mitochondrial morphology and integrity were assessed in SH-SY5Y and ReNcell VM cell lines overexpressing GFP-tagged α-syn (WT/A53T). (A, B) Representative confocal images show MitoTracker Red staining in SH-SY5Y (A) and ReNcell VM (B) cells, revealing qualitative changes in mitochondrial distribution and signal intensity across α-syn variants and overexpression levels. (C, D) Quantitative assessment of mitochondrial integrity was performed by flow cytometry following MitoTracker Red staining in SH-SY5Y (C) and ReNcell VM (D) cells. Data are shown as violin plots and bar graphs representing the mean fluorescence intensity of MitoTracker Red with Dunn’s correction for multiple comparisons. Statistical significance is indicated as p < 0.03 (*), p < 0.002 (**), p < 0.001 (***); non-significant p > 0.12). Column graphs show the mean ± SEM as an alternative representation of staining intensity shifts.

### Real-time functional mitochondrial metabolism analysis reveals cell type-specific metabolic responses to different variants and dose of α-syn

To further investigate the mitochondrial functional consequences of α-syn overexpression in SH-SY5Y and ReNcell VM cells, we performed Seahorse XF Cell Mito Stress Test analysis, which provides dynamic profiling of OCR across multiple mitochondrial parameters (Fig. 5). In SH-SY5Y cells, overexpression of WT α-syn resulted in a marked increase in mitochondrial respiration relative to control cells (Fig. 5A, B). Both basal and maximal OCR values were elevated in α-syn WT low and WT high groups, indicating an adaptive upregulation of oxidative metabolism. The α-syn WT high group demonstrated the highest mitochondrial activity. Nevertheless, this enhanced respiratory profile was not accompanied by significantly elevated spare respiratory capacity. In contrast, SH-SY5Y cells expressing high levels of A53T mutant α-syn displayed a distinct pattern. Both basal and maximal OCR were reduced compared to WT-overexpressing counterparts (Fig. 5A, B). In contrast, ReNcell VM cells exhibited a consistent decline in mitochondrial function across all α-syn overexpressing groups (Fig. 5C, D). WT α-syn overexpression led to a significant reduction in basal OCR. Maximal respiration and spare respiratory capacity were similarly diminished, indicating compromised ability to respond to energetic demands. Notably, ReNcell VM cells expressing A53T mutant α-syn exhibited the most profound deficits, with further reductions across all respiratory parameters, consistent with pronounced mitochondrial dysfunction. Across both WT and A53T variants, basal OCR declined in comparison to control, and spare respiratory capacity was nearly abrogated in high-expressing mutant groups.

**Figure 5.**
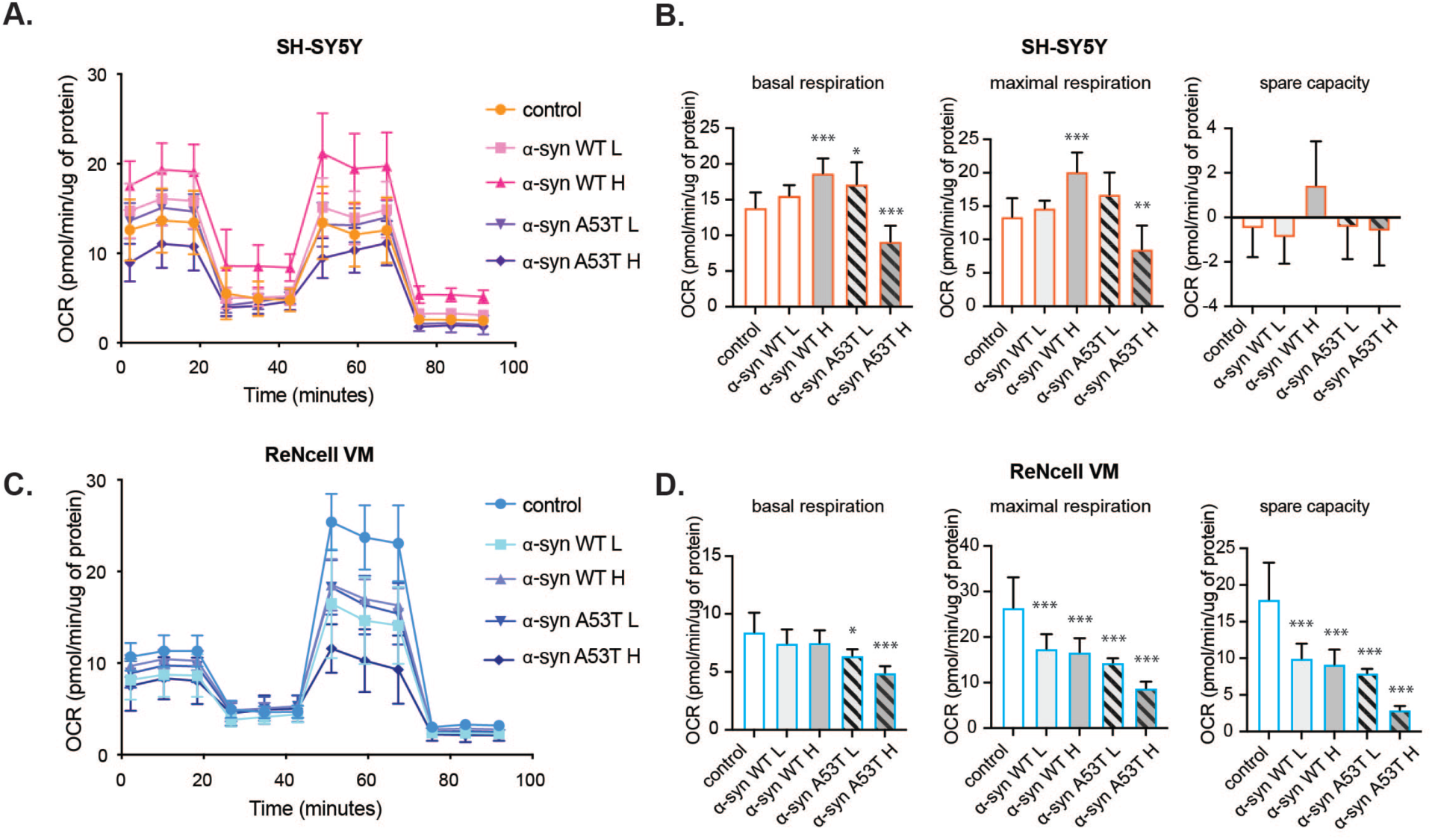
Real-time functional mitochondrial metabolism analysis reveals cell type-specific metabolic responses to different variants and doses of α-syn. Mitochondrial respiration was assessed in SH-SY5Y (A, B) and ReNcell VM (C, D) neuronal cells overexpressing (WT/A53T) α-syn using the Seahorse XF Cell Mito Stress Test. (A, C) Representative OCR traces depict mitochondrial respiratory profiles in SH-SY5Y (A) and ReNcell VM (C) cells following sequential injections of mitochondrial inhibitors and uncouplers. (B, D) Quantification of basal respiration, maximal respiration, and spare respiratory capacity are shown as bar graphs for SH-SY5Y (B) and ReNcell VM (D) cells. Group comparisons were analyzed using one-way ANOVA with Dunnett’s correction for multiple comparisons with significance indicated as p < 0.03 (*), p < 0.002 (**), p < 0.001 (***); non-significant p > 0.12).

### α-syn overexpression and post-translational processing differ between SH-SY5Y and ReNcell VM cell lines

To validate the dosage-dependent overexpression and investigate biochemical processing of α-syn in our cell models, we performed immunoblotting on total protein lysates from SH-SY5Y and ReNcell VM cells stably overexpressing GFP-WT/A53T α-syn (Fig. 6). Syn211 antibody detected a clear, gradual increase in α-syn levels from control to low and high overexpression groups in both SH-SY5Y and ReNcell VM cell lines, confirming the dose-dependent overexpression of α-syn (Fig. 6A, B). In addition to the expected ∼42 kDa band corresponding to GFP-α-syn, a second band migrating approximately 10–12 kDa higher was observed with more prominent bands in WT α-syn high groups. This higher molecular weight band was consistently more prominent in ReNcell VM cells compared to SH-SY5Y cells, suggesting differential processing or PTM of α-syn between the two cell types. ReNcell VM WT α-syn high cells exhibited a significantly higher proportion of the upper band relative to SH-SY5Y WT α-syn high cells and quantitative analysis of the upper to total band ratio further confirmed this difference (Fig. 6C). This cell type-specific difference in band pattern was not observed in the A53T overexpressing lines, where comparable bands were observed across the two cell lines. Probing with a phospho-specific antibody against pS129 revealed increased α-syn phosphorylation with increased overexpression levels in both SH-SY5Y and ReNcell VM cells (Fig. 6A, D). Together, these findings indicate that α-syn overexpression induces distinct biochemical signatures in a cell type-and overexpression level-dependent manner. Based on the difference in band patterns observed especially in the WT α-syn high overexpressing cell lines, we hypothesized that cell type-specific post-translational processing may have a role in the observed band shift. To further investigate this, we immunoprecipitated GFP-α-syn from SH-SY5Y and ReNcell VM cells followed by validation of successful pulldown via immunoblotting (Fig. 6E). Both GFP-WT and GFP-A53T α-syn were efficiently enriched, making it suitable for downstream mass spectrometry analysis (Fig. 7).

**Figure 6.**
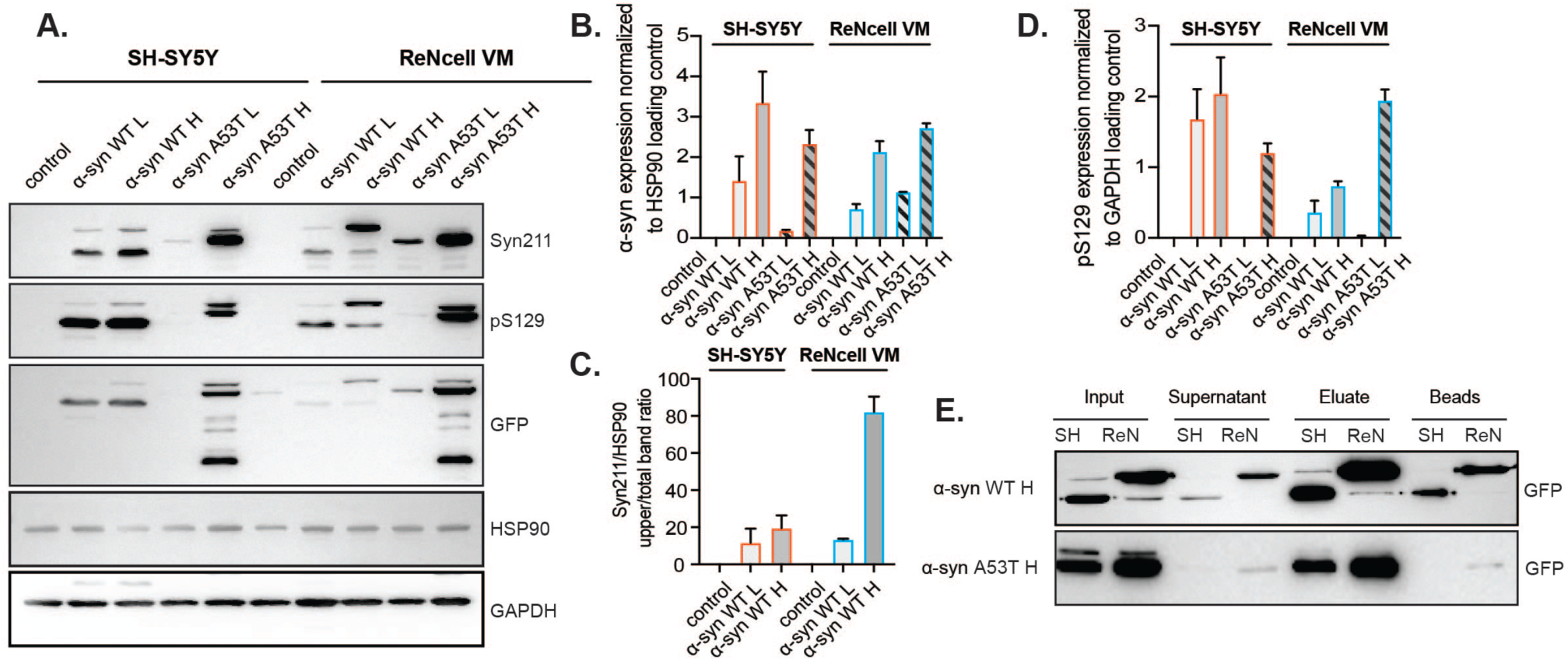
α-syn overexpression and post-translational processing differ between SH-SY5Y and ReNcell VM cell lines. (A) Representative immunoblots showing GFP-tagged α-syn overexpression in SH-SY5Y and ReNcell VM cell lines. Two bands corresponding to GFP-α-syn were detected at approximately 40 kDa and 55 kDa (with Syn211 antibody). Cells with high GFP-α-syn overexpression exhibited markedly increased protein levels compared to low expressers, whereas no signal was observed in untransduced controls. Immunoblot analysis of phosphorylated α-syn at serine 129 (pS129) revealed enhanced phosphorylation in cells with higher GFP-α-syn overexpression. Bands were detected at the same molecular weights as GFP-α-syn. HSP90 (90 kDa) or GAPDH (36 kDa) served as loading control. Quantification of band intensities was performed using ImageJ and normalized to loading controls (HSP90 or GAPDH). Data are presented as mean ± SEM from independent experiments. (B) Total α-syn levels, (C) ratio of upper α-syn band to total α-syn, and (D) levels of phosphorylated α-syn (pS129) are shown. Statistical comparison was not performed for this figure because endogenous α-synuclein levels in the control group were below the detection limit of our assay. As a result, quantifiable measurements were only obtained in the α-synuclein overexpression condition. Since the control values could not be reliably measured, statistical comparisons or multiple group analyses were not applicable. (E) Validation of immunoprecipitation efficiency of GFP-α-syn with elute being subsequently used in mass spectrometry analysis.

**Figure 7.**
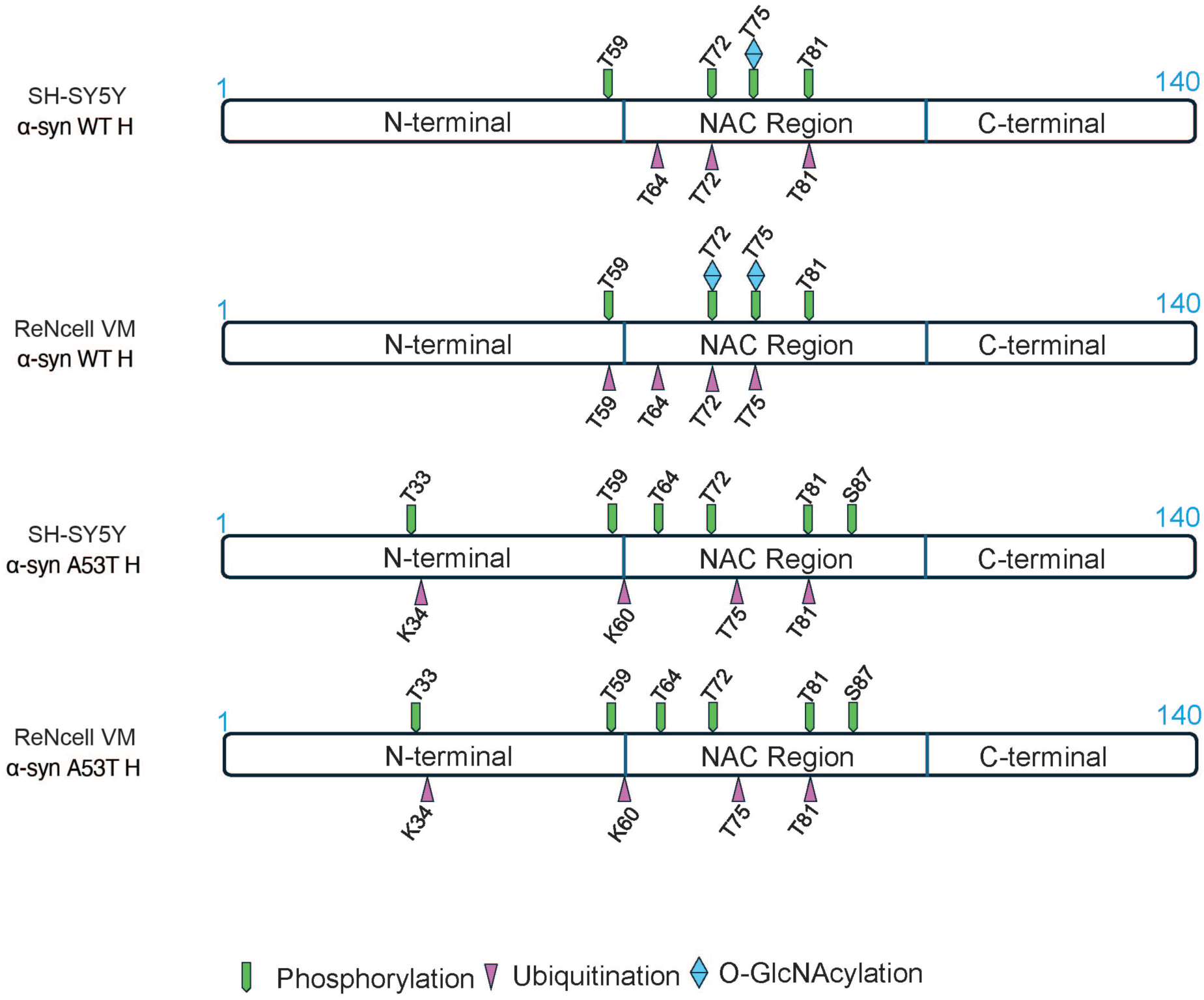
Mass spectrometry identifies NAC region–focused and cell type–specific post-translational modifications of WT but not A53T α-syn. LC/MS analysis was performed on GFP-α-syn immunoprecipitated from SH-SY5Y and ReNcell VM cells. As shown in (A), sequence covered in the analysis was predominantly in NAC region (61-95) and adjacent N-terminal sites. Multiple PTMs including phosphorylation, ubiquitination, and O-GlcNAcylation were identified, with modifications concentrated mainly on threonine residues.

### Mass spectrometry identifies NAC region–focused and cell type–specific post-translational modifications of WT but not A53T α-syn

To investigate the potential molecular basis underlying the distinct α-syn band patterns observed in immunoblotting, we performed mass spectrometry analysis on GFP-tagged α-syn immunoprecipitated from SH-SY5Y and ReNcell VM cells overexpressing either WT or A53T α-syn at high levels. PTMs were mapped across the α-syn sequence. We have selected the PTMs present on residues with peptides detected in both cell types, whether modified or unmodified, to ensure confident cross-sample comparison (Fig. 7). Details of overall abundance and the percentages of modified peptides are provided in the table (Table 1). Sequence coverage was concentrated within the NAC region (61-95) and adjacent N-terminal residues. Within this region, multiple PTMs were identified, including phosphorylation, ubiquitination, and O-GlcNAcylation. Notably, WT α-syn exhibited several cell type–specific differences in modification patterns across six threonine sites. T59 was phosphorylated in both SH-SY5Y and ReNcell VM cells, but was ubiquitinated exclusively in ReNcell VM. T64 was ubiquitinated in both cell lines. T72 showed phosphorylation and ubiquitination in both cell types, while O-GlcNAcylation at this site was only found in ReNcell VM. T75 was phosphorylated and O-GlcNAcylated in both cell lines, but ubiquitination at this residue was again exclusive to ReNcell VM. In contrast, T81 was phosphorylated in both cell types but ubiquitinated only in SH-SY5Y (Fig. 7). These findings reveal a shared phosphorylation profile across backgrounds, but striking divergence in ubiquitination and O-GlcNAcylation, with ReNcell VM cells displaying a broader and more complex PTM profile. The co-occurrence of multiple PTMs at single residues, particularly T72 and T75 suggests potential modification crosstalk and dynamic regulation, which may influence α-syn folding, accumulation, or interactions. Interestingly, we detected ubiquitination on threonine residues, a non-canonical modification rarely reported for α-syn. The presence of threonine ubiquitination across both cell types, with cell type–specific differences in occurrence, indicates the possibility of atypical ubiquitin linkages or non-lysine–based enzymatic targeting under overexpression conditions. In contrast to the variability seen in WT α-syn, the A53T mutant showed identical PTM profiles in both SH-SY5Y and ReNcell VM cells (Fig. 7). All detected residues were modified in the same manner regardless of background, suggesting that the A53T mutation may limit α-syn’s ability to undergo environment-dependent modifications. This loss of PTM plasticity could reflect altered accessibility or reduced interaction with modifying enzymes, potentially contributing to the mutant’s enhanced pathogenicity and reduced clearance observed in disease models. Together, these data reveal that α-syn undergoes diverse, context-dependent PTMs in its NAC and N-terminal regions, with WT protein showing greater sensitivity to the cellular environment than the A53T mutant.

**Table 1.**
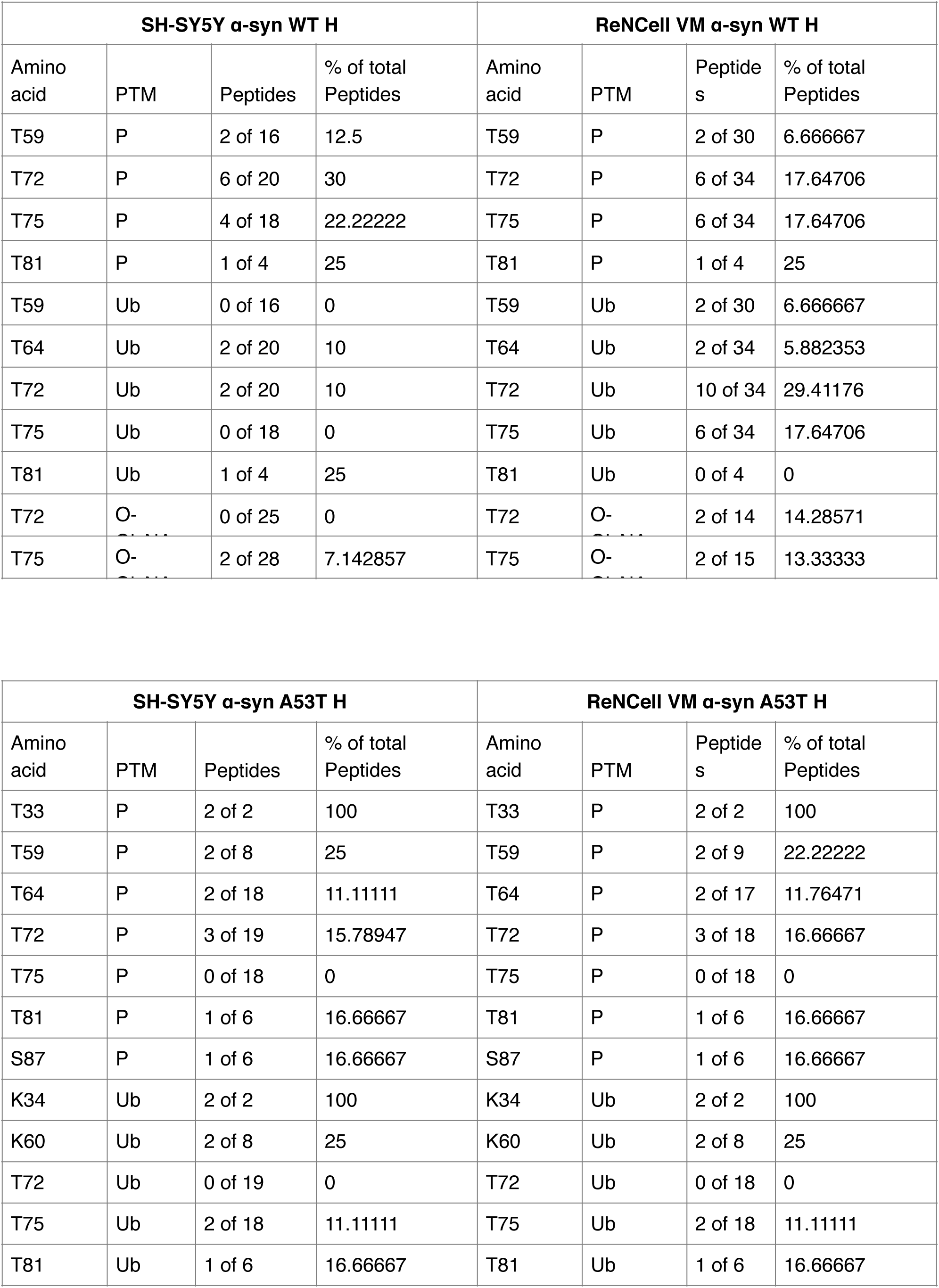
α-syn PTMs detected in LC/MS analysis.

| **SH-SY5Y α-syn WT H** |  |  |  | **ReNCell VM α-syn WT H** |  |  |  |
| --- | --- | --- | --- | --- | --- | --- | --- |
| Amino acid | PTM | Peptides | % of total Peptides | Amino acid | PTM | Peptides | % of total Peptides |
| T59 | P | 2 of 16 | 12.5 | T59 | P | 2 of 30 | 6.666667 |
| T72 | P | 6 of 20 | 30 | T72 | P | 6 of 34 | 17.64706 |
| T75 | P | 4 of 18 | 22.22222 | T75 | P | 6 of 34 | 17.64706 |
| T81 | P | 1 of 4 | 25 | T81 | P | 1 of 4 | 25 |
| T59 | Ub | 0 of 16 | 0 | T59 | Ub | 2 of 30 | 6.666667 |
| T64 | Ub | 2 of 20 | 10 | T64 | Ub | 2 of 34 | 5.882353 |
| T72 | Ub | 2 of 20 | 10 | T72 | Ub | 10 of 34 | 29.41176 |
| T75 | Ub | 0 of 18 | 0 | T75 | Ub | 6 of 34 | 17.64706 |
| T81 | Ub | 1 of 4 | 25 | T81 | Ub | 0 of 4 | 0 |
| T72 | O- | 0 of 25 | 0 | T72 | O- | 2 of 14 | 14.28571 |
| T75 | O- | 2 of 28 | 7.142857 | T75 | O- | 2 of 15 | 13.33333 |

| **SH-SY5Y α-syn A53T H** |  |  |  | **ReNCell VM α-syn A53T H** |  |  |  |
| --- | --- | --- | --- | --- | --- | --- | --- |
| Amino acid | PTM | Peptides | % of total Peptides | Amino acid | PTM | Peptides | % of total Peptides |
| T33 | P | 2 of 2 | 100 | T33 | P | 2 of 2 | 100 |
| T59 | P | 2 of 8 | 25 | T59 | P | 2 of 9 | 22.22222 |
| T64 | P | 2 of 18 | 11.11111 | T64 | P | 2 of 17 | 11.76471 |
| T72 | P | 3 of 19 | 15.78947 | T72 | P | 3 of 18 | 16.66667 |
| T75 | P | 0 of 18 | 0 | T75 | P | 0 of 18 | 0 |
| T81 | P | 1 of 6 | 16.66667 | T81 | P | 1 of 6 | 16.66667 |
| S87 | P | 1 of 6 | 16.66667 | S87 | P | 1 of 6 | 16.66667 |
| K34 | Ub | 2 of 2 | 100 | K34 | Ub | 2 of 2 | 100 |
| K60 | Ub | 2 of 8 | 25 | K60 | Ub | 2 of 8 | 25 |
| T72 | Ub | 0 of 19 | 0 | T72 | Ub | 0 of 18 | 0 |
| T75 | Ub | 2 of 18 | 11.11111 | T75 | Ub | 2 of 18 | 11.11111 |
| T81 | Ub | 1 of 6 | 16.66667 | T81 | Ub | 1 of 6 | 16.66667 |

## Discussion

### Model system design: relevance, stability, and technical advantages

Overexpression-based cellular models have been instrumental in advancing our understanding of α-syn-mediated toxicity in PD. However, the majority of these models rely on inducible expression systems, which can introduce confounding factors such as artificial stress responses and mitochondrial dysfunction. Tetracycline- or doxycycline-based induction, in particular, has been shown to impair mitochondrial protein synthesis, reduce oxygen consumption, and promote mitochondrial fragmentation at commonly used concentrations (0.01–1 µg/mL), thereby compromising experimental outcomes [19–21]. Moreover, doxycycline exposure leads to mito-nuclear protein imbalance and shifts cellular metabolism away from oxidative phosphorylation toward glycolysis, further complicating analyses of mitochondrial and metabolic phenotypes [20, 22, 23]. These well-documented off-target effects limit the interpretability of results in studies where mitochondrial function is a key endpoint. To overcome these limitations, we established a panel of stably expressing cell lines overexpressing either WT or A53T mutant α-syn in two neuronal backgrounds SH-SY5Y neuroblastoma cells and ReNcell VM ventral midbrain progenitor cells. This dual-cell model enabled us to directly compare the effects of α-syn variants, dosage, and cellular context under consistent experimental conditions. Our stable expression system provided several advantages. It maintained consistent α-syn protein levels across low passage numbers (passages 3–5), avoiding variability associated with extended passaging or transient transfection. Although some variability in mRNA levels was observed across biological replicates, protein levels remained stable, consistent with previous reports showing a decoupling between α-syn mRNA and protein expression [24]. This highlights the necessity of directly assessing protein levels when interpreting toxicity or functional outcomes in α-syn models.

Importantly, the expression levels achieved in our system (approximately 3–10-fold above endogenous) fall within the pathophysiological range reported in human PD and synucleinopathies, thereby enhancing the translational relevance of our findings [24]. In contrast to inducible systems, our model minimizes artificial metabolic perturbations, allowing for a clearer interpretation of the intrinsic cellular effects of α-syn.

### Comparative cell biology: cell type-specific vulnerability

#### A53T mutation: pathogenic effects across neural contexts

Our data revealed that the A53T mutation induces remarkably consistent toxic phenotypes across both SH-SY5Y and ReNcell VM cell models, characterized by pronounced ROS elevation, a mutation-driven shift from diffuse to punctate α-syn. High-expression A53T populations also displayed mitochondrial dysfunction, indicating that the A53T variant fundamentally alters α-syn’s mitochondrial interactions. This is in line with studies showing that A53T enhances the ability of α-syn to assemble into oligomers that directly interact with mitochondrial membranes, disrupting normal mitochondrial function and dynamics [25]. The increased MitoTracker Red staining in A53T-overexpressing cells may reflect mitochondrial hyperpolarization resulting from impaired ATP synthase function or altered mitochondrial membrane permeability, representing an early pathogenic event in A53T-mediated neurotoxicity. This convergence aligns with extensive evidence documenting the robust, context-independent pathogenicity of A53T α-syn in neuronal systems [26–28]. The A53T mutation enhances oligomerization of α-syn even at physiological expression levels [25, 29], accelerates fibril formation rates, and has been linked to autosomal dominant early-onset PD with more severe clinical presentations [30]. The mutation fundamentally alters dynamics of α-syn aggregation through enhanced membrane binding [31] and formation of toxic oligomeric intermediates [32, 33]. These oligomers demonstrate region-specific conformational differences, with inclusion-bearing regions harboring oxidized oligomers that accelerate aggregation and cause neuronal degeneration [33]. The consistent pathogenic effects observed across our cellular models support the dominant toxicity of A53T variants, which override cell type-specific protective mechanisms through gain-of-toxic-function mechanisms [32]. Interestingly, despite comparable WT α-syn overexpression levels, only A53T triggered the formation of punctate aggregates. This pattern was independent of expression level and consistently observed across both cell lines, strongly indicating that the A53T mutation possesses an intrinsic aggregation-promoting effect that fundamentally alters conformational behavior of α-syn [32, 33]. This observation aligns with protein engineering studies showing that mutations specifically destabilize native protein states or accelerate conversion of partially folded conformations into oligomeric structures. The A53T mutation enhances oligomer formation and membrane interactions [25, 31], creating toxic species that can disrupt cellular function even at modest expression levels where WT α-syn protein remains diffusely distributed. Importantly, few studies have systematically compared A53T effects across different human neural cell types. Previous comparative work primarily examined SH-SY5Y versus PC12 cells, focusing on autophagy impairment rather than comprehensive metabolic profiling [34]. Our dual-cell system highlights the consistent pathogenic impact of the A53T mutation across distinct neuronal backgrounds, thereby enhancing the relevance and applicability of our findings to broader PD modelling.

#### WT α-syn: neuroprotection and context-sensitivity

The protective effects of WT α-syn overexpression observed in SH-SY5Y cells align with mounting evidence for α-syn’s physiological neuroprotective functions [35]. WT α-syn has been demonstrated to protect against oxidative damage, dopamine toxicity, and various neurotoxic insults through its co-chaperone functions and role in maintaining synaptic vesicle homeostasis [36, 37]. The enhanced mitochondrial function, increased viability, and improved bioenergetic capacity observed in our SH-SY5Y WT α-syn overexpressers likely reflect α-syn’s interaction with ATP synthase, where monomeric α-syn localizes to mitochondria and aids ATP synthase efficiency. The physiological role of monomeric α-syn in mitochondrial bioenergetics is supported by knockout studies where loss of α-syn resulted in uncoupled mitochondrial respiration and reduced ATP production [37, 38]. SH-SY5Y WT α-syn overexpressers showed 40–60% increase in basal respiration, 2–3-fold higher maximal respiratory capacity, and substantially elevated spare respiratory capacity, indicating robust mitochondrial reserve to respond to energetic demands. This mitochondrial enhancement correlates with increased cellular viability and enhanced MitoTracker Red staining, supporting WT α-syn’s potential neuroprotective role through direct interactions with ATP synthase or other mitochondrial components. Additionally, α-syn can rescue neurodegeneration and motor impairment resulting from cysteine-string protein-α deficiency, demonstrating its physiological neuroprotective function at synapses [36]. Our observation of elevated pS129 levels in SH-SY5Y WT high α-syn overexpressers, without associated toxicity, supports recent paradigm-shifting findings that pS129 phosphorylation occurs after initial protein deposition and serves a protective function [39]. This phosphorylation appears to represent a cellular response to manage increased protein burden rather than a pathogenic modification. Multiple lines of evidence demonstrate that pS129 inhibits α-syn fibril formation and seeded aggregation while reducing cellular toxicity [39]. Recent work has established that pS129 is physiologically triggered by neuronal activity to positively regulate synaptic transmission [40, 41], serving as an activity-dependent trigger for protein-protein interactions necessary for the function of α-syn at synapses. In vivo studies demonstrated that the non-phosphorylated form of S129 exacerbates α-syn-induced nigral pathology, whereas S129 phosphorylation eliminates α-syn-induced nigrostriatal degeneration [42]. These findings challenge older paradigms wherein pS129 was viewed solely as a pathological marker.

The contrasting toxicity observed in ReNcell VM cells suggests fundamental differences in cellular stress response mechanisms and highlights the cell type-specific nature of α-syn effects. ReNcell VM cells, derived from ventral midbrain neural progenitors, show proteomic landscapes relevant to dopaminergic neuron development and may better recapitulate the vulnerability of dopaminergic neurons in PD [43]. ReNcell VM cells exhibited consistent mitochondrial dysfunction across all α-syn variants, characterized by reduced basal respiration (20–40% decreases), decreased maximal capacity, and diminished spare respiratory capacity. This pattern of mitochondrial impairment resembles the dysfunction, respiratory collapse, and selective neuronal death observed in PD midbrain dopaminergic neurons [44], highlighting ReNcell VM as a sensitive and disease-relevant model system. The mitochondrial vulnerability in ReNcell VM cells may reflect differences in mitochondrial composition, antioxidant capacity, or protein quality control mechanisms compared to SH-SY5Y cells. This vulnerability pattern corresponds closely to observations in patient-derived neurons and human PD post-mortem brain tissue. The enhanced PTM patterns observed in ReNcell VM WT α-syn high overexpressers, evidenced by altered electrophoretic mobility, may reflect increased ubiquitination at non-canonical sites. Under proteotoxic stress conditions, cells can exhibit promiscuous ubiquitination patterns, potentially including modification at sites like T75 through non-canonical mechanisms involving E2/E3 pairs capable of serine/threonine ubiquitination. This could represent a compensatory response where cells attempt to manage protein overload through alternative degradation pathways, potentially disrupting protective modifications like O-GlcNAcylation at T75.

### Differential post-translational modification landscapes reflect cell-type-specific vulnerability

The distinct PTM profiles observed between SH-SY5Y and ReNcell VM cells offer valuable insight into mechanisms underlying differential neuronal vulnerability. In Western blot analyses, ReNcell VM cells overexpressing WT α-syn at high levels consistently displayed prominent upper molecular weight bands, absent in SH-SY5Y counterparts, suggesting extensive PTMs. These shifts likely reflect a complex modification profile, including enhanced ubiquitination or phosphorylation, representing an adaptive response to increased α-syn burden. Such modifications may serve dual roles: protective, anti-aggregation functions such as O-GlcNAcylation and certain phosphorylation events or as markers of cellular stress that promote protein degradation, such as ubiquitination. The balance between these opposing PTM types plays a critical role in regulating α-syn solubility, clearance, and toxicity. This evolving concept of a context-dependent “PTM code” proposes that cell-specific modification landscapes can dictate whether neurons resist α-syn pathology or enter a pathogenic state. Interestingly, A53T-overexpressing cells did not show comparable differences in PTM profiles between the two cell types. This uniformity further underscores the pathological dominance of the A53T variant and suggests its potential capacity to escape or bypass normal post-translational regulation.

### Clinical relevance and ReNcell VM model validity

The enhanced susceptibility of ReNcell VM cells to α-syn-mediated toxicity may better recapitulate the vulnerability of dopaminergic neurons in PD. ReNcell VM cells, derived from ventral midbrain neural progenitors, represent a reproducible and easy-to-propagate cell culture system for studying neural differentiation with direct relevance to dopaminergic neuron biology [43]. These cells can differentiate into all three neural lineages (neurons, astrocytes, oligodendrocytes) and have been used for PD modeling [43, 45]. Studies using ReNcell VM cells have established functional links between PD-associated genes like PINK1 and neural degeneration, demonstrating their utility as physiologically relevant tools for studying PD mechanisms [45, 46]. The extensive PTMs observed in ReNcell VM cells mirror findings from human post-mortem brain tissue, where complex PTM patterns are associated with synucleinopathy progression [47]. The proteomic landscapes of differentiated ReNcell VM cells show characteristics relevant to dopaminergic neuron development and vulnerability [43]. The differential responses between cell types suggest that therapeutic strategies must consider cellular context. The protective effects observed in SH-SY5Y cells with WT α-syn overexpression support approaches aimed at enhancing α-syn’s physiological functions rather than simply reducing protein levels. Conversely, the vulnerability patterns in ReNcell VM cells suggest that interventions targeting protein quality control mechanisms and PTM pathways may be particularly relevant for vulnerable neuronal populations.

## Conclusion

Our comparative, longitudinal analysis using a stably engineered dual-cell model establishes cell type as a critical determinant of α-syn-driven pathology, independent of simple protein abundance. These findings demonstrate that α-syn’s cellular effects are highly dependent on both genetic context (WT vs A53T) and cellular environment (SH-SY5Y vs ReNcell VM), with the same protein capable of conferring protection or toxicity depending on the specific neuronal context. The consistent pathogenic effects of A53T across cell types support its dominant toxicity. The differential responses to WT α-syn overexpression reveal cell type-specific vulnerability mechanisms that may explain the selective neurodegeneration patterns observed in PD. The enhanced PTMs observed in ReNcell VM cells represent stress-induced compensatory responses that ultimately contribute to cellular dysfunction, highlighting the importance of PTM landscape, mitochondrial adaptation, and cellular quality control in dictating neuronal resilience or vulnerability. Similarly, the protective metabolic effects of WT α-syn in specific cellular contexts support approaches aimed at enhancing α-syn’s physiological functions. These results reinforce the need for targeted, context-aware PD cellular modelling that accounts for neuronal population-specific vulnerabilities and provide a novel platform for further mechanistic and therapeutic investigations.

## Statements and declarations

## Author contributions

Conceptualization, DF; methodology, MM, PM, DM, NM, KA, LF, MD, SS, PP, MB; formal analysis, MM, DF, DM, NM, MB, MKM; investigation, MM, DF; writing – MM original data preparation; writing – review and editing DF, MM, DM; supervision, project administration funding acquisition, DF. All authors read and approved the final version of the manuscript.

## Data availability

All data generated or analyzed during this study are included in this article. The original files and raw mass spectrometry data are available from the corresponding author upon request.

## Ethics approval

This study does not involve any animal, clinical ethics or human participants.

## Abbreviations

α-syn: α-synuclein
FACS: Fluorescence-activated cell sorting
GFP: Green fluorescent protein
OCR: Oxygen consumption rate
PD: Parkinson’s disease
PFA: Paraformaldehyde
PTM: Post-translational modification
ROS: Reactive oxygen species
SEM: Standard error of the mean
WT: Wild-type

